# Min-Redundancy and Max-Relevance Multi-view Feature Selection for Predicting Ovarian Cancer Survival using Multi-omics Data

**DOI:** 10.1101/317982

**Authors:** Yasser EL-Manzalawy, Tsung-Yu Hsieh, Manu Shivakumar, Dokyoon Kim, Vasant Honavar

## Abstract

**Background:** Large-scale collaborative precision medicine initiatives (e.g., The Cancer Genome Atlas (TCGA)) are yielding rich multi-omics data. Integrative analyses of the resulting multi-omics data, such as somatic mutation, copy number alteration (CNA), DNA methylation, miRNA, gene expression, and protein expression, offer the tantalizing possibilities of realizing the potential of precision medicine in cancer prevention, diagnosis, and treatment by substantially improving our understanding of underlying mechanisms as well as the discovery of novel biomarkers for different types of cancers. However, such analyses present a number of challenges, including the heterogeneity of data types, and the extreme high-dimensionality of omics data.

**Methods:** In this study, we propose a novel framework for integrating multi-omics data based on multi-view feature selection, an emerging research problem in machine learning research. We also present a novel multi-view feature selection algorithm, MRMR-mv, which adapts the well-known Min-Redundancy and Maximum-Relevance (MRMR) single-view feature selection algorithm for the multi-view settings.

**Results:** We report results of experiments on the task of building a predictive model of cancer survival from an ovarian cancer multi-omics dataset derived from the TCGA database. Our results suggest that multi-view models for predicting ovarian cancer survival outperform both view-specific models (i.e., models trained and tested using one multi-omics data source) and models based on two baseline data fusion methods.

**Conclusions:** Our results demonstrate the potential of multi-view feature selection in integrative analyses and predictive modeling from multi-omics data.

## Background

The advent of “big data” offers enormous potential for understanding and predicting health risks and intervention outcomes, as well as personalizing treatments, through integrative analysis of clinical, biomedical, behavioral, environmental, and even socio-demographic data. For example, recent efforts in cancer genomics under the Precision Health Initiative offer promising ways to diagnose, prevent, and treat many cancers [1]. Recent advances in high-throughput omics technologies offer cost-effective ways to acquire diverse types of genome-wide multi-omics data. For instance, Large-scale collaborative efforts such as the Cancer Genome Atlas (TCGA) and the International Cancer Genome Consortium (ICGC) are collecting multi-omics data for tumors along with clinical data for the patients. An important goal of these initiatives is to develop comprehensive catalogs of key genomic alterations associated for a large number of cancer types [2, 3].

Computational analyses of multi-omics data offer an unprecedented opportunity to deepen our understanding of complex underlying mechanisms of cancer that is essential for advancing precision oncology (See for example, [4-7]). Because different types of omics data have been shown to complement each other [8], there is a growing interest in effective methods for integrative analyses of multi-omics data [9-11]. The resulting methods have been successfully used to predict molecular abnormalities that affect clinical outcomes and therapeutic targets [5, 10, 12-16]. In general, computational approaches to multi-omics data integration have to address three major challenges [5]: i) the curse of dimensionality (i.e., the number of features *p* is very large compared to the number of samples *n*); ii) the differences in scales as well as sampling/collection bias and noise present in different omics data sets; iii) extracting and optimally combining, for the prediction task at hand, features that provide complementary information across different data sources. Unfortunately, baseline approaches of integrating multiple data sources by simply concatenating their features or analyzing data from each data source separately and combining the predictions fail to satisfactorily address these challenges. Therefore, there is an urgent need for more sophisticated methods for integrative analysis and predictive modeling from multi-omics data [16].

Multi-view learning offers a promising approach to developing predictive models by leveraging complementary information provided by multiple data sources (views) [17]. Multi-view learning algorithms attempt to learn one model from each view while jointly optimize all the view-specific models to improve the generalization performance. Some examples of multi-view learning algorithms include: multi-view support vector machines [18], multi-view Boosting [19], multi-view *k*-means [20], and clustering via canonical correlation analysis [21]. The problem of learning predictive models from multi-omics data can be naturally formulated as a multi-view learning problem where each omics data source constitutes a view. However, the vast majority of existing multi-view learning algorithms are not equipped to effectively cope with the high-dimensionality of omics data [22]. Hence, predictive modeling from multi-omics data calls for effective methods for multi-view feature selection or dimensionality reduction.

Against this background, we describe an approach to predictive modeling from multi-omics data based on a novel multi-view feature selection algorithm and a two-stage framework for multi-omics data integration. To the best of our knowledge, this is the first attempt to apply multi-view feature selection in the integrative analyses of multi-omics data. We evaluated our approach on the task of predicting ovarian cancer survival [13] using a TCGA multi-omics dataset composed of three omics data sources, copy number alteration (CNA), DNA methylation, and gene expression RNA-Seq. Our results show that: (i) the predictive models developed from multi-omics data (multiple views) outperform their single view counterparts; and that (ii) the predictive models developed using the proposed multi-view feature selection algorithm outperform those developed using two baseline methods that combine multiple views into a single view. These results demonstrate the viability of multi-view feature selection for multi-omics data integration and lay the ground for developing effective multi-omics data integration models using multi-view feature selection.

## Methods

### Datasets

Normalized and preprocessed multi-omics ovarian cancer datasets (most recently updated on August 16, 2016), including gene-level copy number alteration (CNA), DNA methylation, and gene expression (GE) RNA-Seq data, were downloaded from UCSC Xena cancer genomic browser [23]. Table 1 summarizes the number of samples and features (e.g., genes) in each dataset. Clinical data about vital status and survival for the subjects were also downloaded from Xena server. Only the patients with CNA, methylation, RNA-Seq, and survival data were retained. Patients with survival time ≥ 3 years were labeled as long-term survivors while patients with survival time < 3 years and vital status of 0 were labeled as short-term survivors. The resulting multi-view dataset consists of 215 samples, 127 of them are classified as long-term survivors. Each view was then pre-filtered and normalized as follows: i) features with missing values were excluded; ii) feature values in each sample were rescaled to lie in the interval [0, 1]; iii) features with variance less than 0.02 were removed.

**Table 1.** TCGA ovarian cancer omics data used in this study

| Data source | Platform | Number of samples | Number of features | No of features with high variance |
| --- | --- | --- | --- | --- |
| CNA | Affymetrix SNP 6 | 579 | 24,777 | 7,355 |
| Methylation | Illumina Infinium HumanMethylation27k | 616 | 27,579 | 6,206 |
| GE RNA-Seq | Illumina HiSeq | 308 | 30,531 | 283 |

### Notations

Table 2 summarizes convenient notations used in this work. For simplicity, we assumed a binary label for each sample. Note however, that Algorithms 1 and 2, described below, are also applicable to multi-class as well as numerically labeled data.

**Table 2.** Notations

| Symbol | Definition and Description |
| --- | --- |
| $D = \langle X, y \rangle$ | Labeled dataset where $X \in R^{m \times n}$ is a matrix of $m$ instances and $n$ features, and $y \in \{0,1\}^m$ is the binary class labels of the instances |
| $x_i$ | $i^{\text{th}}$ feature in $X$ |
| $g(x_i, x_j)$ | Function that returns the redundancy between two features $x_i$ and $x_j$ |
| $f(x_i, y)$ | Function that returns the relevance between a feature $x_i$ and class labels $y$ |
| $S$ | Indices of selected features |
| $\Omega$ | Indices of all features |
| $\Omega_S$ | Indices of candidate features $\Omega - S$ |
| $k$ | Number of features to be selected |
| $v$ | Number of views in a multi-view dataset |
| $MVD = \langle (X^1, \dots, X^v), y \rangle$ | Labeled multi-view dataset where $X^i \in R^{m \times n_i}$ is a matrix of $m$ samples and $n_i$ features and $y \in \{0,1\}^m$ is the binary class labels of the instances in all views |
| $D^i = \langle X^i, y \rangle$ | $i^{\text{th}}$ view in a multi-view dataset |
| $x_j^i$ | $j^{\text{th}}$ feature in $X^i$ |
| $S^i$ | Indices of selected features from $i^{\text{th}}$ view |
| $\Omega^i$ | Indices of all features in $i^{\text{th}}$ view |
| $\Omega_{S^i}$ | Indices of candidate features $\Omega^i - S^i$ in $i^{\text{th}}$ view |

### Minimum redundancy and maximum relevance feature selection

Unlike univariate feature selection methods [24] that return a subset of features without accounting for redundancy between the selected features, the minimum redundancy and maximum relevance (MRMR) based feature selection algorithm [25] iteratively selects features that are *maximally relevant* for the prediction task and *minimally redundant* with the set of already selected features. MRMR has been successfully used for feature selection in a number of applications including microarray gene expression data analysis [25, 26], prediction of protein sub-cellular localization [27], epileptic seizure [28], and protein-protein interaction [29].

While the exact solution to the problem of MRMR selection of *k* = |*S*| features from a set of *n* candidates requires O(*n^k^*) searches, it is possible to obtain an approximate solution using a simple heuristic algorithm (see Algorithm 1) [25]. Algorithm 1 accepts as input: a labeled dataset *D*; a function *g*: (*x_i_, x_j_*) → *R*^+^ that quantifies the redundancy between any pair of features (e.g., the absolute value of Pearson’s correlation coefficient); a function *f*: (*x_i_, y*) → *R*^+^ that quantifies the relevance of a target feature for predicting the labels *y* (e.g., mutual information or F-statistic); and the number of features *k* to be selected using the MRMR criterion. In lines 1 and 2, the algorithm creates an empty set *S* and the feature with the maximum relevance for predicting *y* is added to *S*. In each of the subsequent *k* – 1 iterations (lines 3-5), the features that greedily approximate the MRMR criterion in Eq. 1 are successively added to *S*. Eq. 1 has two terms: the first term maximizes the relevance condition, whereas the second term minimizes the redundancy condition.

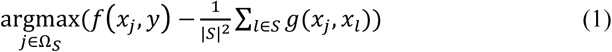

~~~
**Algorithm 1.** MRMR
**Require**: *D* =< *X*,*y* >,*g*, *f*, *k*
1: Φ → S
2: add 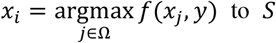 to *S*
3: **for** *t* = 1: *k* − 1 do
4:  add the index of the feature that satisfies Eq. 1 to *S*
5: **end for**
6: **return** S
~~~

### Multi-view minimum redundancy and maximum relevance feature selection

MRMR, or any single-view feature selection algorithm, can be trivially applied to multi-view data as follows to yield two baseline methods: i) Apply MRMR separately to each view and then concatenate view-specific selected features. The major limitation of this approach is that it ignores the redundancy and complementarity of features across views [30]; ii) Apply MRMR to a single-view dataset obtained by concatenating all the views. A key limitation of this approach is that it fails to explicitly account for the prediction task specific differences in the relative utility or relevance of the features extracted from the different views.

We propose a novel multi-view feature selection algorithm, MRMR-mv, an adaptation of MRMR, to the multi-view setting. The proposed MRMR-mv algorithm is shown in Algorithm 2. MRMR-mv accepts as input: a labeled multi-view dataset, *MVD*, with *ν* ≥ 2 views; a redundancy function *g*; a relevance function *f*; number of features to be selected *k*; and a probability distribution *P* = {*p*_1_…*p_ν_*} that models the relative importance of each view (or the prior probability that a view contributes a feature to the set of features selected by MRMR-mv). If all of the views are considered equally important, *P* should be a uniform distribution. MRMR-mv proceeds as follows. First, *S*^*t*^ is initialized for each view *t* to keep track of selected features from that view (lines 1-3). Second, the procedure *choice*, implemented in NumPy python library [31], is invoked, to sample with replacement, *k* − 1 times, according to the selection probability distribution *P* and the list of sampled views is kept in 𝐶 (lines 4 and 5). Third, the maximally relevant feature across all of the views is retrieved, and added to *S*^*i*^, where *i* is the view index of the retrieved feature 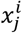 (line 6). Fourth, at each iteration *t*, the feature from the sampled view 𝐶[*t*] that satisfies the MRMR criterion with respect to the previously selected features from all the views examined during the preceding iterations is added to *S*^𝐶[*t*]^ (lines 7-10). Finally, the algorithm returns selected view-specific feature indices, *S*.

We note that MRMR-mv is different from the first baseline approach of separately applying MRMR to each view since MRMR-mv expands the minimum redundancy condition to include selected features from all views while in the former approach redundancy is determined using view-specific features only. Unlike the second baseline approach of applying MRMR to concatenated views, MRMR-mv enables views to be treated unequally (using user’s beliefs about the relative importance of the views for a given prediction task) and jointly performs feature selection in a view-aware manner that allows features from all views to be present in the set of selected features.

### A two-stage feature selection framework for integrating multi-omics data

Figure 1 shows our proposed two-stage framework for integrating multi-omics data for virtually any prediction task (e.g., predicting cancer survival and predicting clinical outcome). The input to our framework is a labeled multi-view dataset in the form *D^i^* =< *X^i^*, *y* >. Stage I includes view-specific filters that can be used to encapsulate any traditional single-view feature selection method (e.g., Lasso [32] or MRMR). Each filter has a gating signal that could be used to disable that filter in which case the disabled filter passes on no data to the 2^nd^ stage. A special view-specific filter, called AllFilter, passes the data from *all* of the input features without performing any feature selection. Stage II has a single filter that can encapsulate either a single-view or multi-view feature selection algorithm. If the 2^nd^ stage filter encapsulates a single-view feature selection method, the feature selection method will be applied to the concatenation of the Stage II input. On the other hand, if the 2^nd^ stage filter encapsulates a multi-view feature selection method (e.g., MRMR-mv), then the multi-view feature selection method will be applied to the multi-view input of Stage II. The framework supports two modes of operations: i) training mode, where each enabled filter will be trained using the input so as to produce the filtered version of the input; ii) test (or operation) mode, where test multi-view dataset is provided as input and the trained filters will output the selected features of the input data.

**Figure 1.**
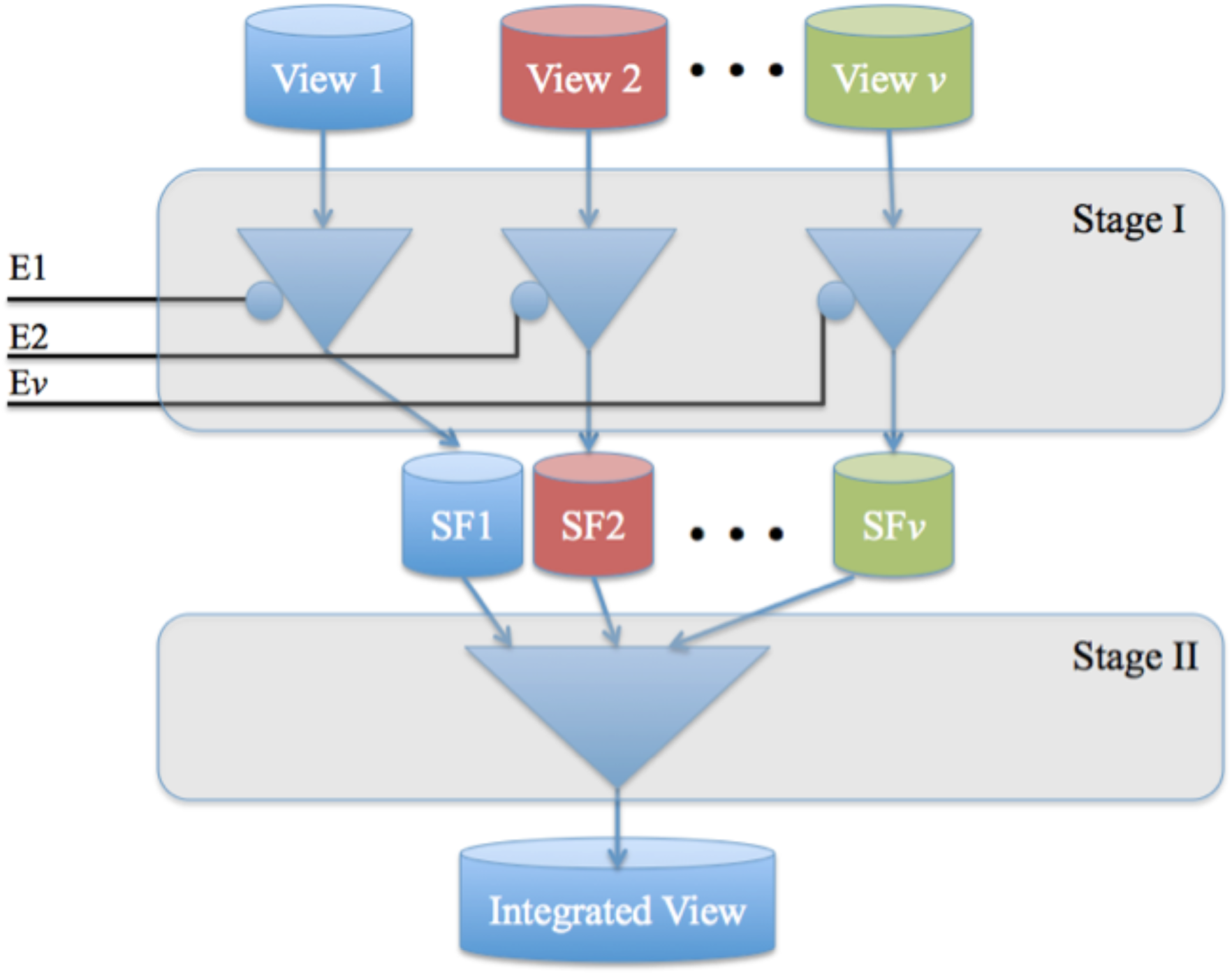
Two-stage framework for integrating multi-omics data. Ei refers to the enable signal for the *i*^th^ view-specific filter. SF*i* refers to the set of features selected from the *i*^th^ view using the *i*^th^ filter.

~~~
**Algorithm 2.** MRMR-mv
**Require:** *MVD* = <(*X*^1^,…,*X^ν^*), *y* >, *g, f, k, P* = (*p_i_,* …,*p_ν_*)
1: for *t* = 1: *ν*
2:    *S*^*t*^ ← Φ
3: end for
4: *V* ← {1, …,*ν*}
5: 𝐶 ← *choice*(*V*, *k* – 1, *P*)
6: add 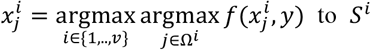
7: **for** *t* = 1:*k* − 1 do
8:    *l* ← 𝐶[*t*]
9:    add 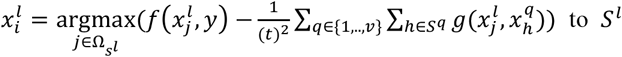
10: **end for**
11: **ruturn** *S* = {*S*^1^,…, *S*^*ν*^}
~~~

The framework can be easily customized so as to allow evaluation of different approaches of predictive modeling from multi-omics data. For example, to build a single view model by applying the Lasso method to the *i*^th^ view, we: set *E_i_* to 1 and disable all other filters; pass Lasso feature selection method to the *i*^th^ filter; use AllFilter as Stage II filter. Similarly, to apply MRMR to concatenated views, we: enable Stage I filters and use either AllFilter (to pass the input as is) or any single-view filter; and deploy MRMR as the Stage II filter.

## Implementation

We implemented Algorithms 1 and 2 and the two-stage feature selection framework in Python using the scikit-learn machine learning library [33]. We will release the code as part of sklearn-fuse, a python library for data and model-based data fusion that is currently under development. In the mean time, the code for the methods described above will be available to interested researchers upon request.

## Experiments

We report results of experiments on the task of building a predictive model of cancer survival from an ovarian cancer multi-omics dataset derived from the TCGA database. The resulting data set is comprised of three views, namely, CNA, methylation, and gene expression RNA-Seq for each patient along with the corresponding clinical outcomes (short-term versus long-term survival). Our first set of experiments consider single-view classifiers based on each of the 3 views to obtain view-specific models for comparison with the proposed multi-view models; The second set of experiments compare some of the representative instantiations of the two-stage multi-view feature selection framework in combination with some representative choices of (single-view) supervised algorithms for training the classifiers. In both cases, we experimented with three widely used machine learning algorithms for developing cancer survival predictors: i) Random Forest (RF) [34] with 500 trees; ii) eXtreme Gradient Boosting (XGB) [35] with 500 weak learners; ii) Logistic Regression (LR) [36] with L1 regularization. We used the implementations of these algorithms available in the Scikit-learn machine learning library [33].

For Stage I feature selection, we experimented with several feature selection methods implemented in Scikit-learn including: RF feature importance [34]; Lasso [32]; ElasticNet [37]; and Recursive Feature Elimination (RFE) [38]. However, due to space limitation, we describe only the results of the best performing method, Lasso with L1 regularization parameter set to 0.0001. In Stage II feature selection, we used MRMR as a baseline method and MRMR-mv for multi-view feature selection

For both MRMR and MRMR-mv feature selection, we used the absolute value of Pearson’s correlation coefficient as the redundancy function, *g*. For the relevance function, *f*, we experimented with three functions Chi2, F-Statistic (F-Stat), and Mutual Information (MI). All functions are implemented in Scikit-learn.

We estimated the performance of the resulting classifiers on the task of predicting cancer survival using the 5-fold cross-validation (CV) procedure. Briefly, the dataset is randomly partitioned into five equal subsets. Four of the five subsets are collectively used to select the features and train the classifier and the remaining subset is held out for estimating the performance of the trained classifier. This procedure is repeated 5 times, by setting aside a different subset of the data for estimating model performance. The 5 results from all the folds are then averaged to report a single performance estimate. In our experiments we used the area under ROC curve (AUC) [39] to assess the predictive performance of classifiers. With small size (in terms of the number of samples) datasets, the estimated classifier performance might vary for different random partitioning of the data into 5 folds (see Section 3.1 for details). To obtain a more robust estimate of performance, we ran the 5-fold cross-validation procedure 10 times (each using different partitioning of the data into 5 subsets) and reported the average AUC estimated from the 10 5-fold CV experiments.

## Results and Discussion

### Single-view models for predicting ovarian cancer survival

We evaluated RF, XGB, and LR classifiers trained using each of the individual views with the top *k* features selected using Lasso feature selection algorithm for choices of *k* = 10,20,30, …,100. Tables 3-5 report the performance of the resulting classifiers averaged over 10 different 5-fold cross-validation experiments.

**Table 3.** Average AUC scores of RF, XGB, and LR models estimated using 10 runs of 5-fold cross validation experiments and CNA data.

| # Features | RF | XGB | LR |
| --- | --- | --- | --- |
| 10 | 0.57 | 0.56 | 0.58 |
| 20 | 0.61 | 0.61 | 0.61 |
| 30 | 0.61 | 0.61 | 0.61 |
| 40 | 0.63 | 0.62 | 0.61 |
| 50 | 0.64 | 0.64 | 0.62 |
| 60 | 0.65 | 0.65 | 0.63 |
| 70 | 0.65 | 0.65 | 0.63 |
| 80 | 0.65 | 0.65 | 0.62 |
| 90 | 0.66 | 0.66 | 0.63 |
| 100 | 0.66 | 0.66 | 0.62 |
| Max | 0.66 | 0.66 | 0.63 |
| Avg. | 0.63 | 0.63 | 0.62 |

**Table 4.**
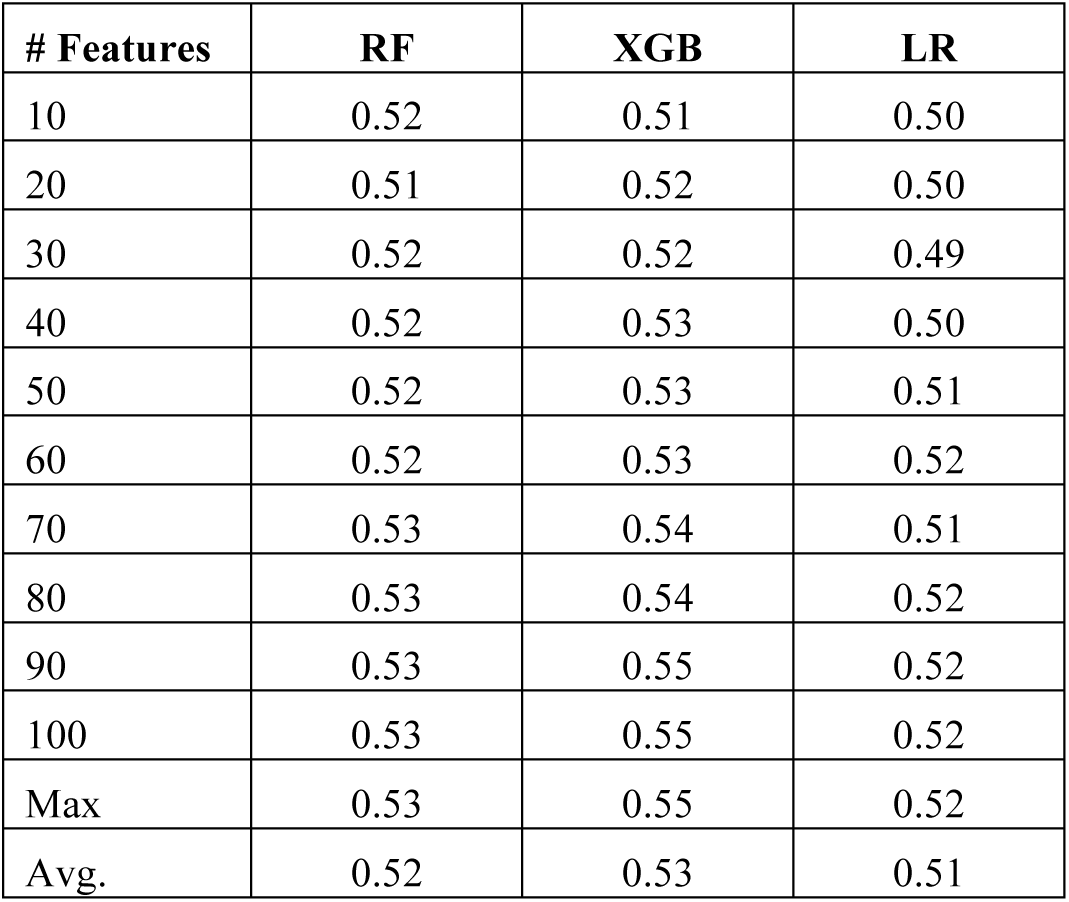
Average AUC scores of RF, XGB, and LR models estimated using 10 runs of 5-fold cross validation experiments and methylation data.

**Table 5.** Average AUC scores of RF, XGB, and LR models estimated using 10 runs of 5-fold cross validation experiments and RNA-Seq data.

| # Features | RF | XGB | LR |
| --- | --- | --- | --- |
| 10 | 0.58 | 0.57 | 0.59 |
| 20 | 0.60 | 0.58 | 0.61 |
| 30 | 0.61 | 0.60 | 0.63 |
| 40 | 0.62 | 0.61 | 0.64 |
| 50 | 0.62 | 0.61 | 0.65 |
| 60 | 0.63 | 0.60 | 0.66 |
| 70 | 0.63 | 0.60 | 0.64 |
| 80 | 0.64 | 0.60 | 0.65 |
| 90 | 0.63 | 0.61 | 0.65 |
| 100 | 0.64 | 0.61 | 0.65 |
| Max | 0.64 | 0.61 | 0.66 |
| Avg. | 0.62 | 0.60 | 0.64 |

We observed that models built using only the methylation view performed marginally better than random guessing (i.e., the best observed average AUC in Table 5 is 0.55). In contrast, single view models using CNA or RNA-Seq achieved higher average AUC scores of up to 0.66. These results are in agreement with those of previously reported studies (e.g., [13]). It should be noted that when the performance of single view models is estimated using a single 5-fold cross-validation experiment (as opposed to average over 10 different cross-validation experiments), the best observed AUC scores were 0.70, 0.55, and 0.69 for models built from the CNA, methylation, and RNA-Seq views, respectively. The observed variability in performance among different 5-fold cross-validation experiments is expected because of the relatively small size of the ovarian cancer survival dataset. This finding underscores the importance of using multiple CV experiments to obtain robust estimates and comparisons of classifier performance. Next, we show how integrating data sources (i.e., views) can further improve the predictive performance of the cancer survival predictors.

### Integrative analyses of multi-omics data sources using multi-view feature selection

We used our two-stage feature selection framework (See Figure 1) to construct multi-view models (MV) with the following settings. The input to the framework is two views, CNA and RNA-Seq. We chose not to use the methylation view because the performance of single view models built using the methylation data performed marginally better than chance (see Section 3.1). For the Stage I filters, we used Lasso with L1 regularization parameter set to 0.0001 to select the top 100 features from CNA and RNA-Seq views, respectively. For the Stage II filter, we used MRMR-mv with Pearson’s correlation coefficient as the redundancy function and a uniform distribution for the selection probability parameter, *P*. Finally, we experimented with different multi-view models obtained using combinations of choices for the remaining MRMR-mv parameters, *k* and *f*. Specifically, we experimented with *k* = 10, 20, …, 100 and the relevance function *f* ∈ {*Chi*2, *F* – *Stat*, *MI, and CFM*} where *CFM* is the average of the other three relevance functions.

Figure 2 compares the performance of the different MV models described above. Interestingly, no single relevance function consistently outperforms other functions for different choices of the number of selected features, *k*, and machine learning algorithms. However, the best AUC of 0.7 is obtained using either *Chi*2 or *MI* relevance functions and RF classifier trained using the top 100 features. Hence, our final MV models will use *Chi*2 as the relevance function and the remaining MRMR-mv settings stated in the preceding paragraph.

**Figure 2.**
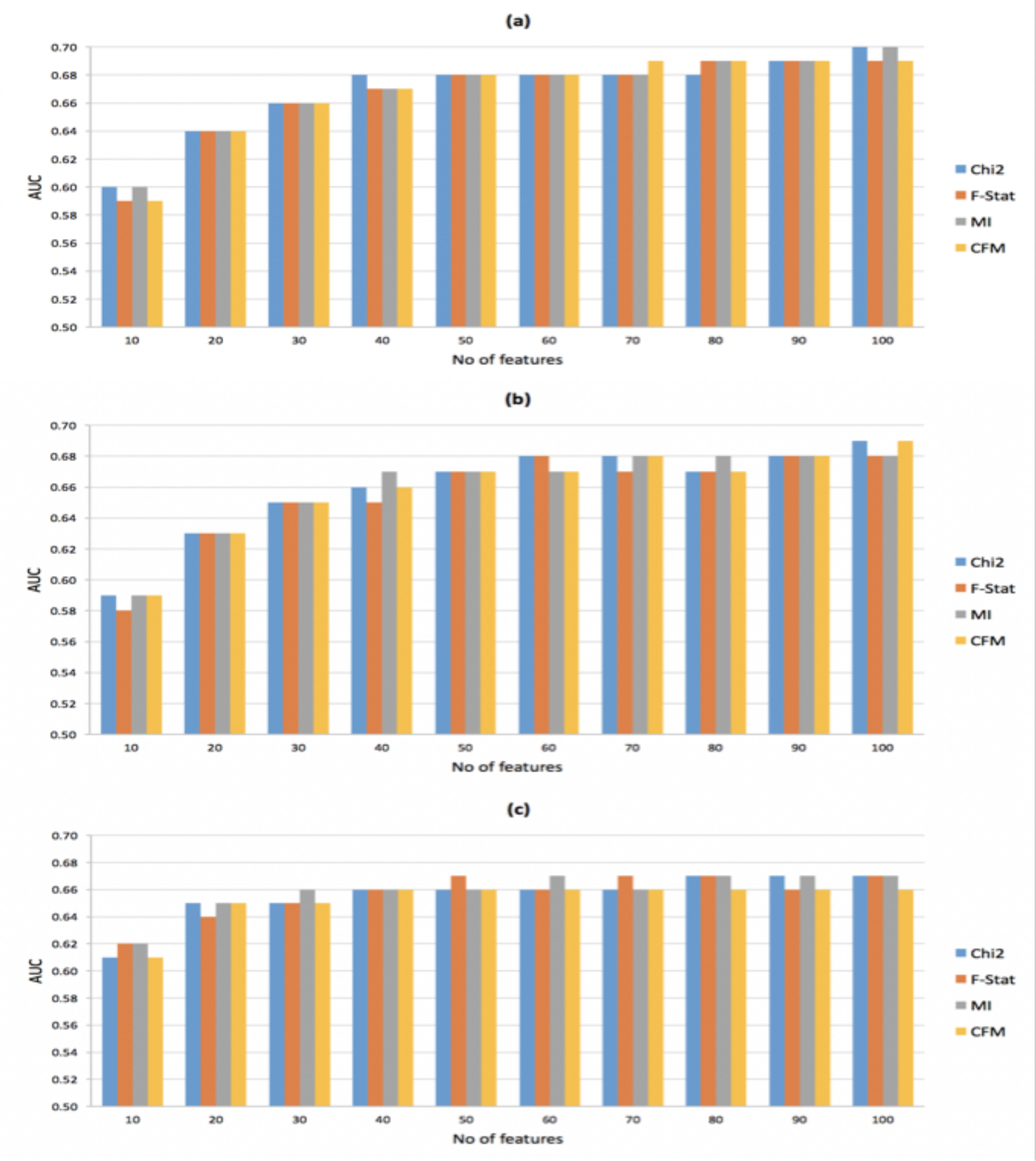
Performance comparisons of multi-view models using four different relevance functions for MRMR-mv and three machine learning classifiers, a) RF, b). XGB, and c) LR

The selection probability parameter, *P*, in MRMR-mv algorithm controls the expected number of selected features from each view. Results shown in Figure 2 have been produced using a uniform selection probability distribution. Although using a uniform distribution is reasonable since the best AUC score for the single-view models based on CNA or RNA-Seq is 0.66 (See Tables 3 and 5), it is interesting to examine the influence of *P* on the performance of our MV models. Let *P* = (*p*_1_, *p*_2_)be the probability distribution where *p*_1_ and *p*_2_ denotes the sampling probability for CNA and RNA-Seq, respectively. In this experiment, we considered 11 different probability distributions obtained using *p*_1_ = {0,0.1, 0.2, …,1}. Then, for each choice of the number of selected features, *k*, we evaluated 11 MV models using RF algorithm and the same MRMR-mv settings described in the preceding subsection and the 11 different probability distributions for *P*. We used the percent relative range in the recorded AUC to assess the sensitivity of MV models to changes in *P*. Figure 3 shows the relationship between the number of selected MV features, *k*, and the sensitivity of MV models to changes in *P*. Interestingly, our results suggest that as the number of selected MV features increases, the resulting MV models become less sensitive to the selection probability distribution parameter *P*.

**Figure 3.**
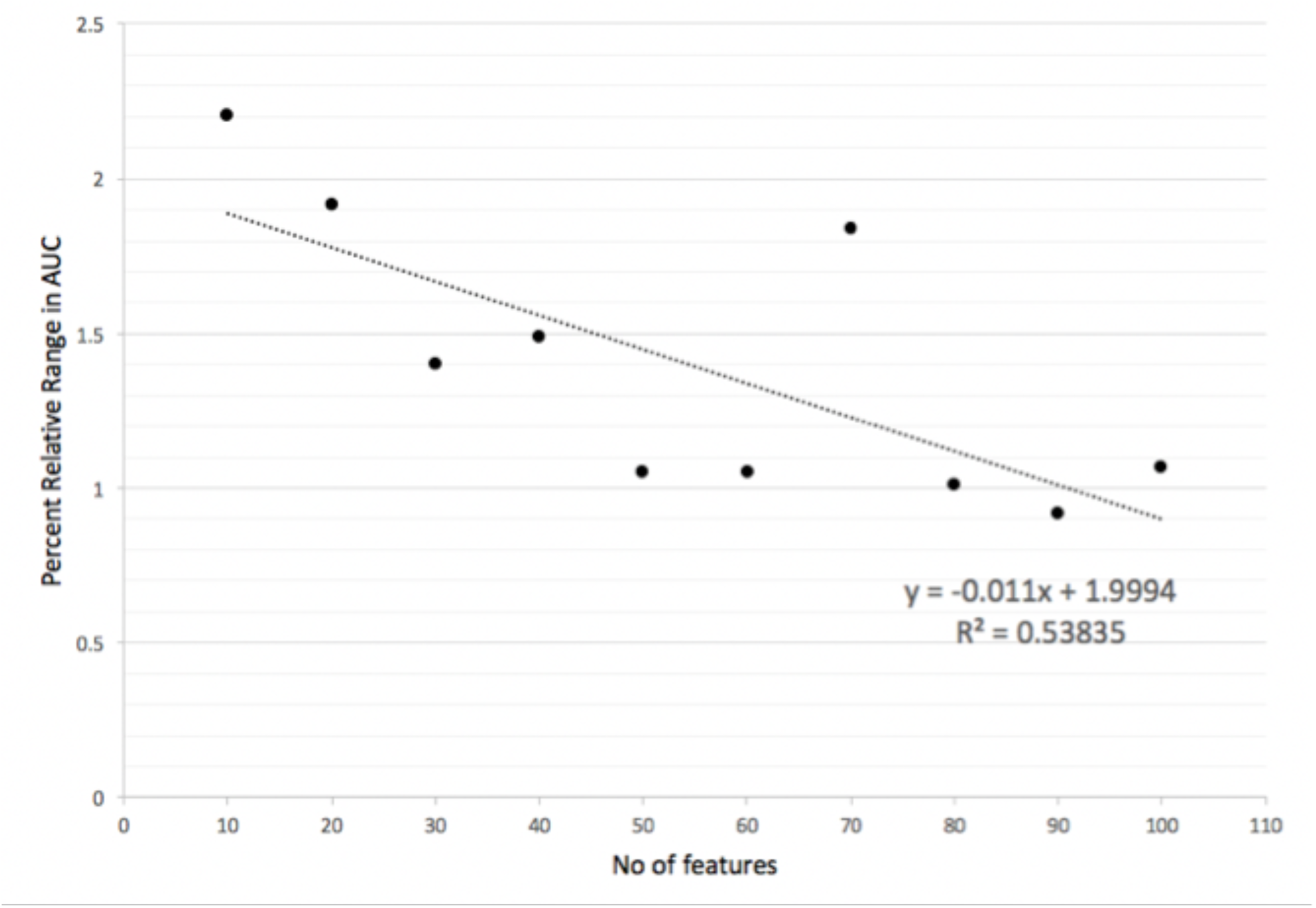
Relationship between No of selected MV features and sensitivity of MV models to changes in selection probability distribution P in terms of percent relative range in AUC.

### Multi-view vs. single-view models for predicting ovarian cancer survival

Figure 4 compares our final MV models with the following single-view models: i) SV_CNA, single-view models developed using CNA data source; ii) SV_RNA-Seq, single-view models developed using RNA-Seq data source; iii) SV_C, single-view models obtained by applying MRMR to the *concatenation* of the two views, CNA and RNA-Seq; iv) SV_S, single-view models obtained by applying MRMR *separately* to CNA and RNA-Seq views, respectively. In addition, Figure 4 shows the results for a simple ensemble model that averages the predictions from SV_CNA and MV models. In general, MV and Ensemble models outperform SV models in most of the cases.

**Figure 4.**
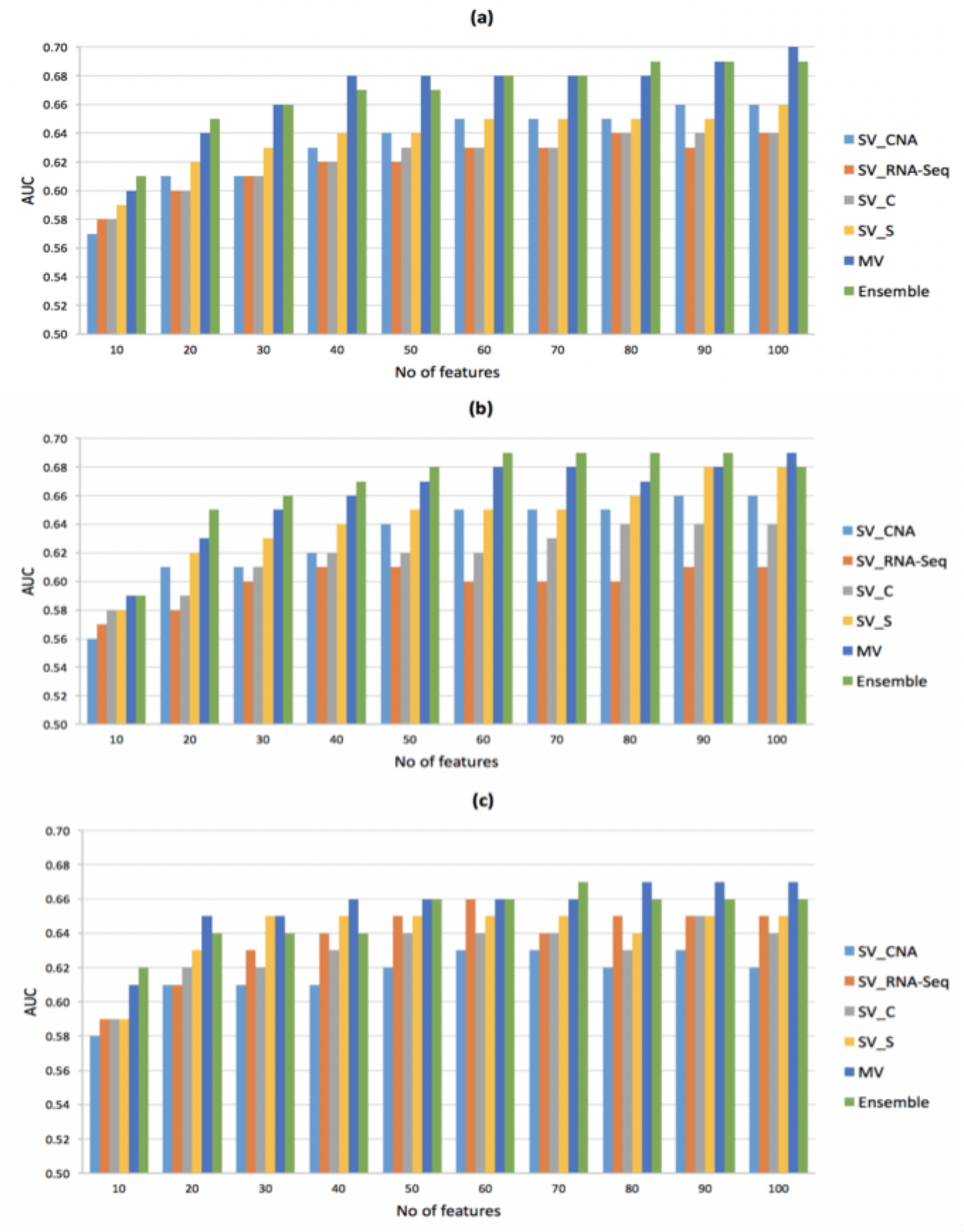
Performance comparisons of final multi-view models with single-view models using three machine learning classifiers, a) RF, b) XGB, and c) LR.

We noted some interesting observations from our experiments with each of the machine learning algorithms considered in our experiments. In the case of models developed using RF algorithm, MV and Ensemble models outperformed the four single-view models for all choices of the number of selected features, k. Ensemble models outperformed MV models for *k* = 10, 20, and 80. Baseline single-view models outperformed SV_CNA and SV_RNA-Seq for *k* ≤ 40. The highest observed AUC was 0.7 and was obtained using the MV model and *k* =100. In the case of XGB based models, SV_S, MV, and Ensemble models outperformed the remaining single-view models. Ensemble models outperformed MV models for 8 out of 10 choices of k. Finally, for models developed using LR algorithm, SV_S, MV, and Ensemble models outperformed the other three single-view models. Regardless of which machine learning algorithm was used, SV_RNA-Seq and SV_C models had the lowest AUC in most of the cases reported in Figure 4. Our results suggest that the best single-view model is more likely to perform better than models developed using concatenated views. Our results also suggest that either applying feature selection to each individual view or selecting features jointly using multi-view feature selection consistently outperform the best single view model.

### Analysis of the top selected multi-view features

In order to get insights into the most discriminative features selected by our framework, we considered the top 100 features selected using MRMR-mv jointly from CNA and RNA-Seq views. To determine which features (genes) could serve as potential biomarkers for ovarian cancer survival, at each of the 50 iterations (resulting from running 5-fold procedure for 10 times), we scored each per-view input feature (input to our framework) by how many time it appears in the top 100 features. Table 6 summarizes the top 20 features from each view along with their normalized feature importance scores.

**Table 6.**
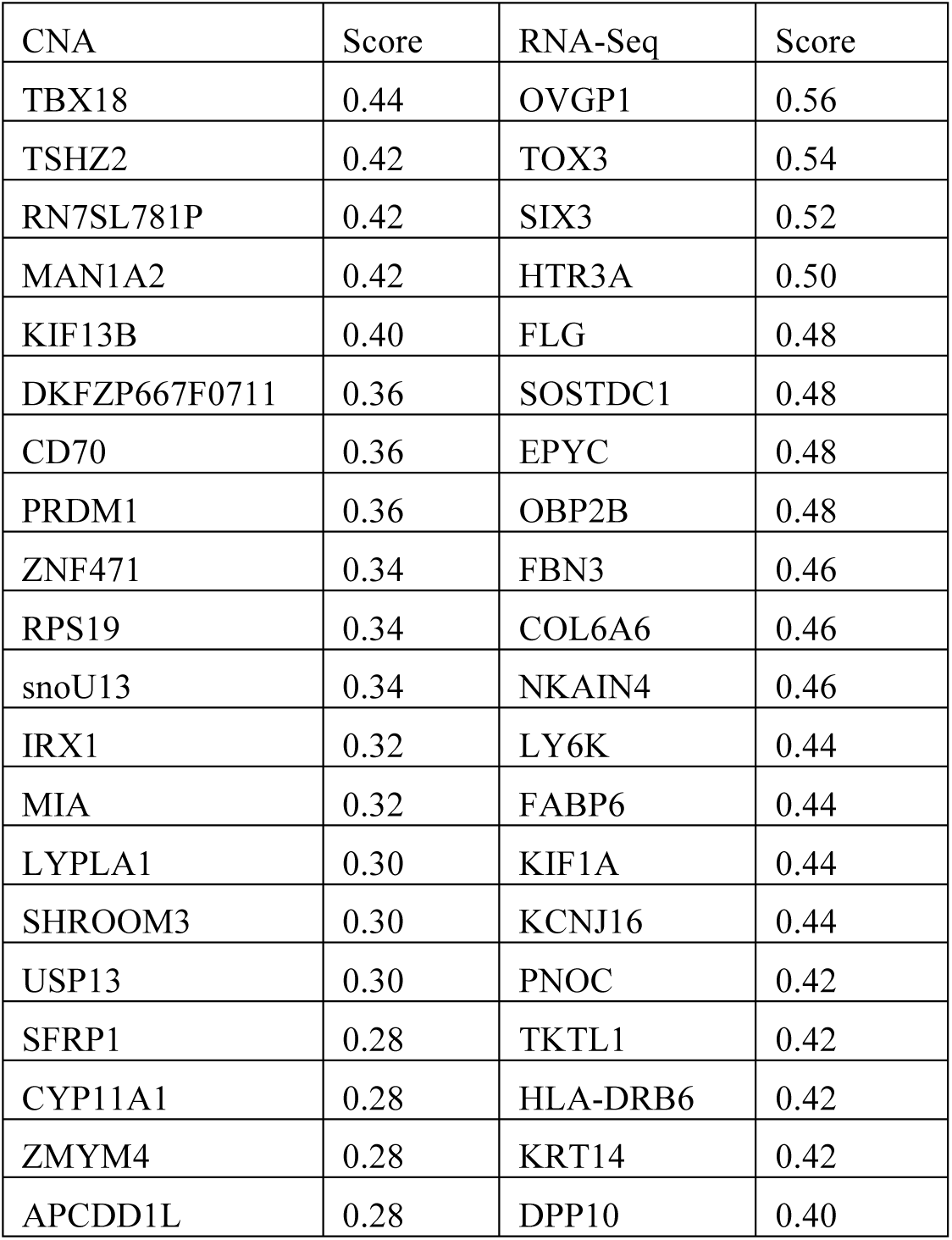
List of top 20 selected features from CNA and RNA-Seq views

To examine the interplay between the top selected features from each view, we constructed an integrated network of interactions among the features using the cBio portal by integrating the biological interactions from public databases including NCI-Nature Pathway Interaction Database, Reactome, HPRD, Pathway Commons, and MSKCC Cancer Call Map [40]. Examination of the resulting network (Figure 5) shows that *RPS19, PNOC*, *SFRP1* and *KCNJ16* are connected to other frequently altered genes, including *MYC or EIF3E* as oncogenes, from TCGA ovarian cancer dataset. In particular, ribosomal protein S19 (*RPS19*), which is known to be up-regulated in human ovarian and breast cancer cells and released from apoptotic tumor cells, was found to be associated with a novel immunosuppressive property [41]. Furthermore, *HTR3A* is targeted by several FDA approved cancer drugs retrieved from PiHelper [42], an open source compilation of drug-target and antibody-target associations derived from several public data sources.

**Figure 5.**
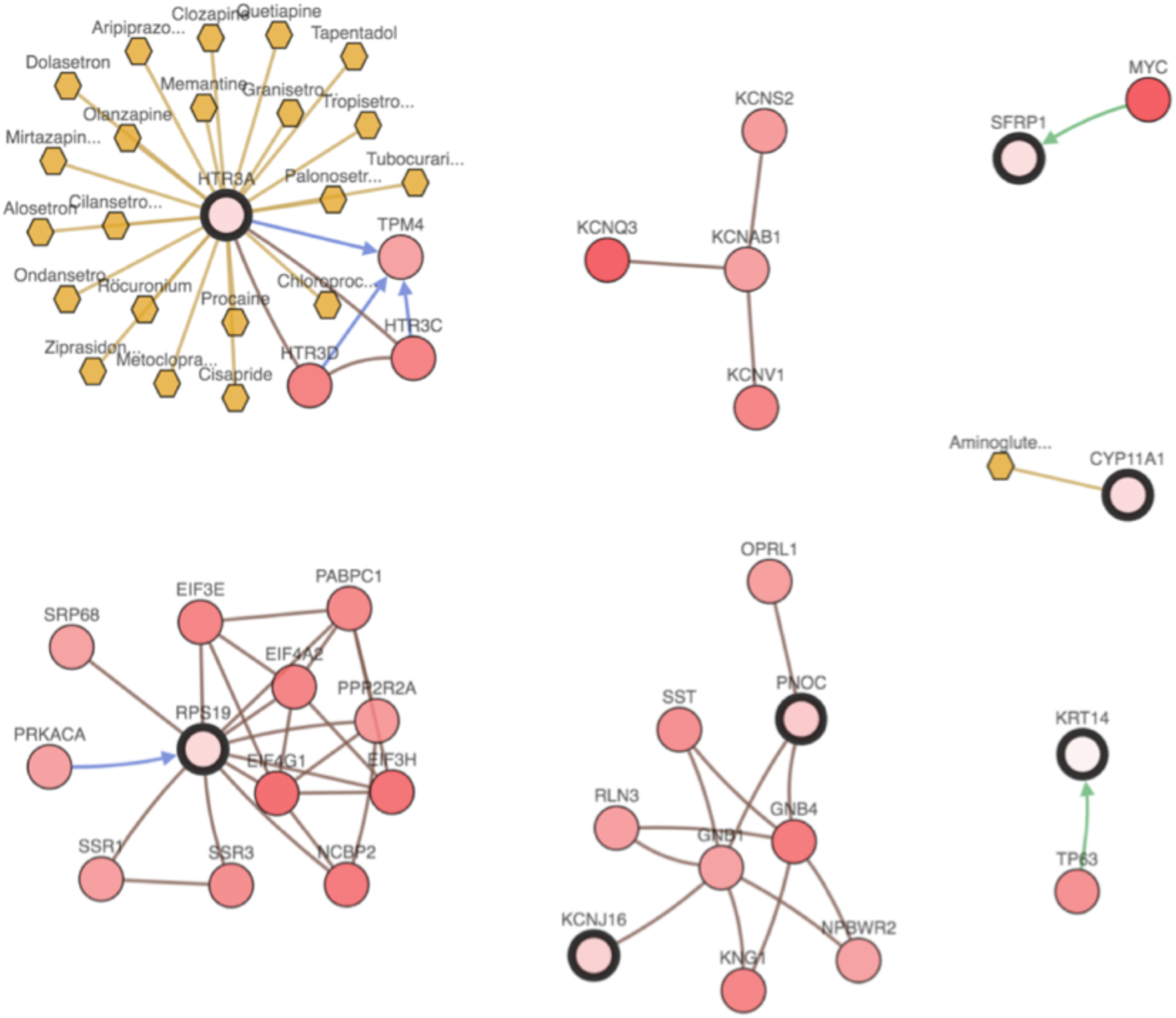
Integrative network view of selected features from CNA and RNA-Seq views. Selected features are highlighted by a thicker black outline. The remaining nodes are frequently altered neighbor genes with known interactions with the highlighted genes and were derived from public interaction databases. Each node in the network is gradient color-coded according to the alteration frequency based on CNA and RNA-Seq data derived from the TCGA ovarian cancer dataset via cBio Portal.

Finally, we performed a gene-set enrichment analysis to identify overrepresented GO terms in the two sets of top 20 features from CNA and RNA-Seq views. Specifically, we used the gene-batch tool in GOEAST (Gene Ontology Enrichment Analysis Software Toolkit) [43] with default parameters to import the gene symbols and to identify significantly overrepresented GO terms, for Biological Processes, Cellular Components and Molecular Function categories, in the CNA and RNA-Seq features sets. We found that the selected CNA gene set was enriched with 220 GO terms whereas the selected RNA-Seq gene set was enriched with 40 GO terms (See additional files 1 and 2). Analysis of the GO terms enriched in the CNA gene set showed a significant over-representation of the molecular function GO terms related to hydrolase activity, oxidoreductase activity, and ion binding. Analysis of the GO terms enriched in the RNA-Seq gene set showed a significant over-representation of the molecular function GO terms related to transmembrane and substrate-specific transporter activity. We also used the Multi-GOEAST tool to compare the results of enrichment analysis of CNA and RNA-Seq gene sets. The graphical outputs of the Multi-GOEAST analysis results for top selected genes in CNA and RNA-Seq in Biological Processes, Cellular Components and Molecular Function categories are provided in additional files 3-5. In these graphs, red and green boxes represent enriched GO terms only found in CNA and RNA-Seq, respectively. Yellow boxes represent commonly enriched GO terms in both sets of genes. The saturation degrees of all colors represent the significance of enrichment for corresponding GO terms. Interestingly, GO:0003777~microtubule motor activity term is only shared GO term between CNA and RNA-Seq enriched terms (see additional file 5). We concluded that the CNA and RNA-Seq features selected by the proposed multi-view feature selection algorithm are non-redundant not only in terms of the genes selected from the CNA and RNA-Seq views but also in terms of their significantly overrepresented GO terms.

### Conclusions

Developing multi-omics data-driven machine learning models for predicting clinical outcome, including cancer survival, is a promising cost-effective computational approach [44]. However, the heterogeneity and extreme high-dimensionality of omics data present significant methodological challenges in applying the state-of-the art machine learning algorithms to training such models from multi-omics data. In this paper, we have described, to the best of our knowledge, the first attempt at at applying multi-view feature selection to address these challenges. We have introduced a two-stage feature selection framework that can be easily customized to instantiate a variety of approaches to integrative analyses and predictive modeling from multi-omics data. We have proposed MRMR-mv, a novel maximum relevance and minimum redundancy based multi-view feature selection algorithm. We have applied the resulting framework and algorithm to build predictive models for ovarian cancer survival using multi-omics data derived from the Cancer Genome Atlas (TCGA). We have demonstrated the potential of integrative analysis and predictive modeling of multi-view data in ovarian cancer survival prediction. Work in progress is aimed at further development of multi-view feature selection and predictive modeling methodologies and their application in translational biomedical data sciences.

## Acknowledgments

We gratefully acknowledge the TCGA Consortium and all its members for the TCGA Project initiative, for providing sample, tissues, data processing and making data and results available. The results published here are in whole or part based upon data generated by The Cancer Genome Atlas pilot project established by the NCI and NHGRI. Information about TCGA and the investigators and institutions that constitute the TCGA research network can be found at http://cancergenome.nih.gov.

## Availability of data and materials

The processed TCGA datasets used for analysis are publicly available at https://xenabrowser.net/datapages/

## Authors’ contributions

YE, DK, and VH designed and conceived the research. YE developed and implemented the feature selection algorithm and the data integration framework. YE and TH ran the experiments. YE, TH, and MS performed the data analysis. YE drafted the manuscript. DK and VH edited the manuscript. All authors read and approved he final version of the manuscript.

## Ethics approval and consent to participate

Not applicable.

## Consent for publication

Not applicable.

## Supplementary Material

**Additional file 1**: GOEAST gene-batch output of enriched GO terms in the Biological Processes, Cellular Components and Molecular Function categories for CNA top selected genes

**Additional file 2**: GOEAST gene-batch output of enriched GO terms in the Biological Processes, Cellular Components and Molecular Function categories for RNA-Seq top selected genes

**Additional file 3**: Graphical output of Multi-GOEAST analysis results of Biological Processes GO terms in the top selected genes in CNA and RNA-Seq.

**Additional file 4**: Graphical output of Multi-GOEAST analysis results of Cellular Components GO terms in the top selected genes in CNA and RNA-Seq

**Additional file 5**: Graphical output of Multi-GOEAST analysis results of Molecular Function GO terms in the top selected genes in CNA and RNA-Seq

